# Abortion occurs during double fertilisation and ovule development in *Paeonia ludlowii*

**DOI:** 10.1101/2021.07.16.452677

**Authors:** Ting-qiao Chen, Meng-yu Xie, Yu-meng Jiang, Tao Yuan

## Abstract

*Paeonia ludlowii* (Stern & G.Taylor) D.Y.Hong, a rare and endangered species, is indigenous to Tibet, China and propagated only by seed. Its natural reproduction is constrained by low fecundity. Excess seed abortion is a key factor restricting its natural reproduction, cultivation, introduction, and protection. In this study, we examined the characteristics of aborted ovules, developmental differences after flowering of normal and aborted ovules, and their ratios at different positions in *P. ludlowii* ovary. During pollination, fertilisation, and seed development, ovule abortion was frequent, with a random abortion position. There were four types of abortion, namely, abnormal pistil, sterile ovules, abnormal embryo sac, embryo and endosperm abortions. Of these, embryo and endosperm abortions could be divided into early abortion, middle abortion, and late abortion. The early aborted ovules stopped growing on day 12, the endoblast and endosperm in the embryo sac aborted gradually. Furthermore, the shape of the embryo sac cavity changed. The volume of aborted ovules was significantly different from that of fertile ovules. At ripening, the external morphology of different types of aborted seeds was significantly different.

**Summary statement:** Elucidation of the origin and characteristics of ovule abortion in an endangered Chinese plant species, *Paeonia ludlowii*.

## Introduction

*Paeonia ludlowii* (Stern & G.Taylor) D.Y.Hong, *Paeonia* Sect. *Moutan* DC., has high ornamental, economic, breeding, medicinal, and development values (Fig. 1) (Hong, 1997; Li et al., 2018; Li et al., 2019; Lu et al., 2020; Zhang et al., 2020; Lu et al., 2021). Until the end of the 20^th^ century and the beginning of the 21^st^ century, *P. ludlowii* was listed as an independent species (Hong, 1997; Hong & Pan, 2005). Wild *P. ludlowii* was only distributed in a small area in the eastern Himalayas, Tibet, China; to date, only six wild populations have been reported (Hong et al., 2017). All known *P. ludlowii* species originate from these regions (Cheng et al., 1998).This species has been introduced in several areas in China, but it does not flourish in all areas; in fact, in only four areas, the species could flower and bear fruits normally (Li, 2005; Ni, 2009; Cui et al., 2019).

**Fig. 1.**
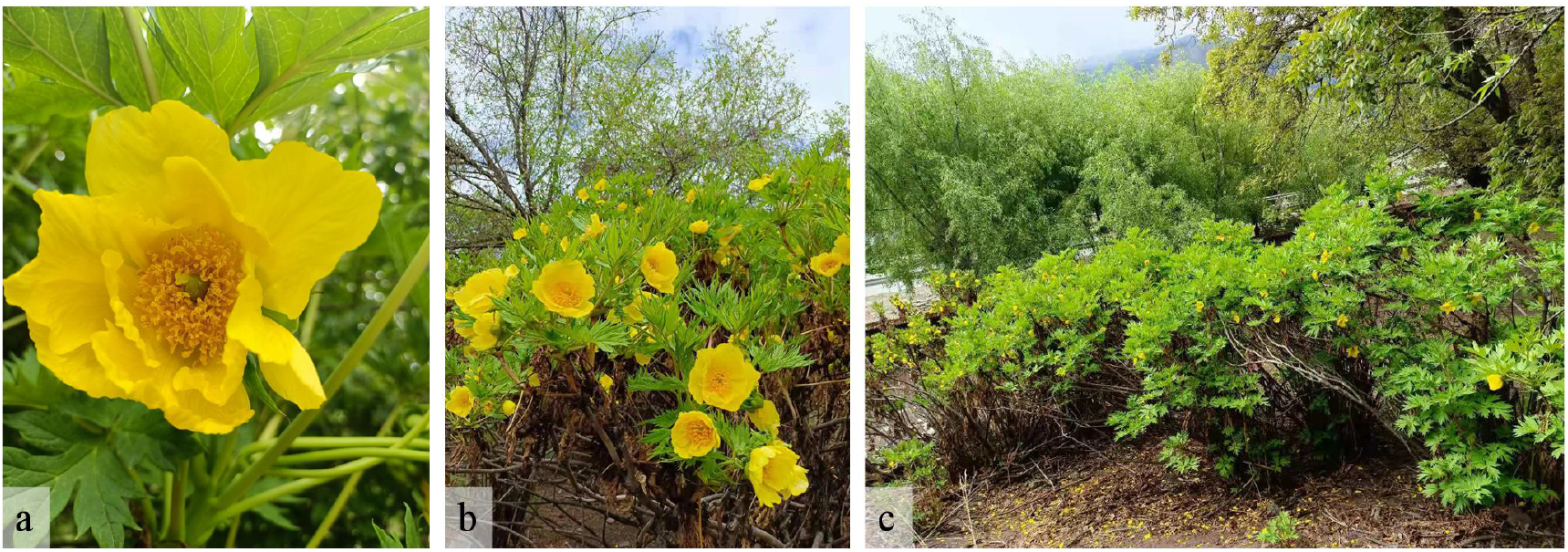
Paeonia ludlowii: photo taken in Nyingchi, Tibet on May 10, 2021

*Paeonia ludlowii* can only be propagated from the seeds (Cheng et al., 1997), and hence, it needs to produce a large number of seeds and germinate easily in order to expand its population. However, in the wild, the rate of seed setting and sprouting of P. ludlowii is extremely low, and the seeds have an extremely long dormancy period (Yang et al., 2007; Ma et al., 2012; Hao et al., 2014). Moreover, due to over-excavation, catastrophes, and severe anthropogenic environmental destruction, wild *P. ludlowii* is endemic to China and is listed as ‘critically endangered’ in the Endangered Species Red Book of China (Wang and Xie, 2004). Therefore, it is necessary to study seed abortion in *P. ludlowii*.

Seed abortion is common in most flowering plants and is considered a potentially beneficial mechanism for improving the quality of offspring (Burd, 1998;Arathi et al., 1999;Miyajima et al.,2003; Satoki et al., 2009; Meyer et al., 2014; Oneal et al., 2016; Li et al., 2021). However, the mechanism of abortion severely restricts the proliferation, breeding, introduction, and conservation of endangered species. In fact, the phenomenon of ovule abortion in tree peony (*P. suffruticosa*) is common. The abortion rate of wild *P. jishanensis* could be as high as 85% (Luo et al., 1998), and that of *P. rockii* is 50% (Cheng, 1996). Existing research shows that the amount of pollen with normal development is substantial and that it could meet the demand of all ovules in each carpel. Moreover, most of the ovules of tree peony develop normally (Cheng et al., 1999;Pan et al., 1999), but the surface structure of the stigma is primitive and simple, which is not conducive to pollination (Zhao, 2002). Hence, it has been suggested that abnormal pollination, fertilisation, and seed development might lead to ovule abortion in tree peony.

A seed is the result of ovule development, and hence, to investigate seed abortion, ovule development must first be fully understood. Therefore, in this study, fluorescence microscopy and scanning electron microscopy (SEM) were used to observe the processes of pollen tube growth and fertilisation to elucidate sexual reproduction in *P. ludlowii* after pollination. We also selected paraffin sections to explore the internal structure of the embryo sac before and after ovule fertilisation to determine the differences between fertile and aborted ovules and the specific origin and timing of aborted ovules. We aimed to compare fertile and abortive ovules, by examining the developmental characteristics of abortive ovules and the number and positions of abortive ovules in a mature ovary. We evaluated the occurrence and developmental characteristics of abortive ovules. This study is of significance to reproductive biology research on *P. ludlowii.* It provides primary data regarding the proliferation and breeding of *P. ludlowii* and lays a theoretical foundation for further studies on seed abortion.

## Results

### Normal ovule fertilisation and seed development

#### Morphological characteristics of the carpel and embryo sac

The carpel of *P. ludlowii* consists of three parts: stigma, style, and ovary (Fig. 2 a). The plant has a wet stigma, and it is curled in various forms, mostly about 360°. It is formed by combining two parts of similar size and shape, and a narrow band of width 0.1–0.4 mm is formed in the middle (Fig. 2 b). The surface of the stigma is densely covered with papillary cells (Fig. 2 c). The style is joined to the stigma, approximately 1–3-mm long, with a hollow style canal in the centre (Fig. 2 e). The style canal is the growth channel for the pollen tube of *P. ludlowii.* After entry into the style, the pollen tubes grow downward along the hollow style canal and the inner canal cells of the style. The nuclei of these cells are large and obvious; morphologically, the cells are regular and darker than other cells. They have the characteristics of glandular cells (Fig. 2 f). Attached to the style is the ovary, in which are two rows of inverted ovules in a linear arrangement (Fig. 2 d), with a double integument, pellicle, and thick nucellus (Fig. 2 g). At blooming, the embryo sac is mature and forms a typical seven-cell and eight-nucleus structure (Fig. 2 h–k; Fig. 2 h, two synergids; Fig. 2 i, an egg cell; Fig. 2 j, three antipodal cells; Fig. 2 k, central cells [two polar nuclei]). Subsequently, the two polar nuclei fuse to form a secondary nucleus (Fig. 2l); but occasionally, they do not fuse to form a secondary nucleus. Only one embryo sac was observed in each ovule in all experimental materials.

**Fig. 2.**
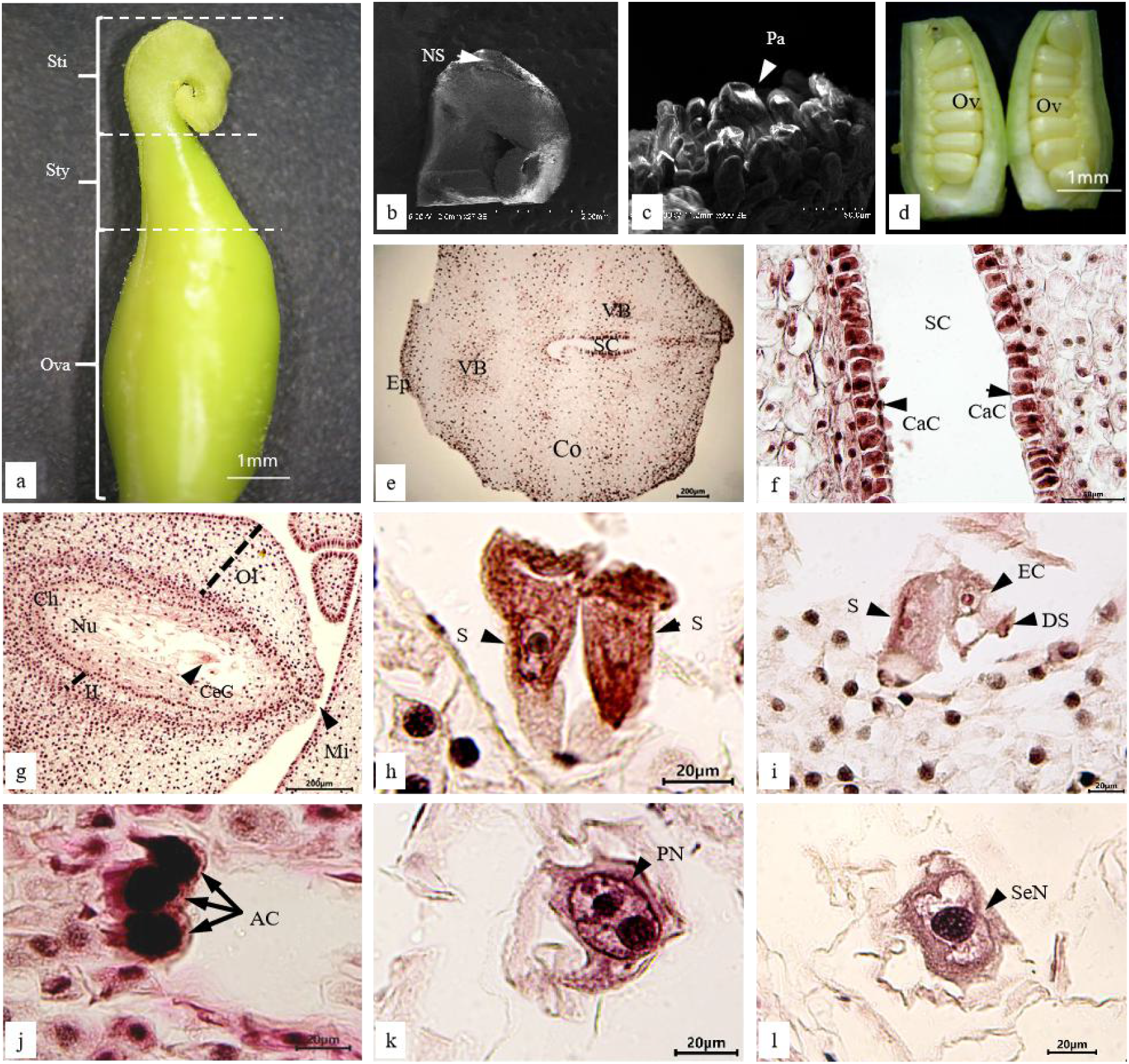
Carpel and embryo sac of *P. ludlowii:* **(a)** the carpellate form of *P. ludlowii,* consisting of the stigma, style, and ovary; **(b)** stigma curving nearly 360° and is formed by two parts of similar size and shape, and a narrow strip is formed in the middle (NS); **(c)** the papillary cells on the surface of the stigma; **(d)** two rows of ovules in the ovary; **(e)** transverse section of the style, a hollow stylistic tract can be seen; **(f)** longitudinal section of style, the large and distinct inner epidermal cells of the style can be clearly seen; **(g)** longitudinal section of an ovule showing the internal structure of the ovule; **(h)** the two synergids were located at the micropyle end, the nuclei were obvious, and there were obvious filamentous organelles at the micropyle end; **(i)** after flowering for 12 h, in the egg, one synergid had degenerated; **(j)** the material inside the antipodal cells is intensively stained; **(k)** a central cell having two polar nuclei; and **(l)** the central cell in which the two polar nuclei fuse to form the secondary nucleus. AC, antipodal cell; CeC, central cell; CaC, canal cell; Ch, chalaza; Co, cortex; DS, degenerated synergid; EC, egg cell; Ep, epidermis; II, inner integument; Mi, micropyle; Nu, nucellus; NS, narrow strip; OI, outer integument; Ov, ovule; Ova, ovary; Pa, papilla; PN, polar nucleus; S, synergid; Sti, stigma; Sty, style; SC, style canal; VB, vascular bundle; SeN, secondary nucleus.

#### Pollen tube growth in the style and ovary

The flowers of *P. ludlowii* open during the day and close at night, a feature that facilitates pollination by wind and insects. We observed that 1–12 h after flowering, pollen grains fell on the stigma and germinated, the pollen tube penetrated deep into the stigma, and the pollen remained on the stigma (Fig. 3 a). After entering the stigma, the pollen tubes grew along the vascular bundles of the stigma and converged into the style (Fig. 3 b). The tube then grew to the base through the stylistic tract and the layer of mucous secreted by the inner epidermal gland cells (Fig. 3 c). After 36–48 h of flowering, the pollen tube reached the base of the style (Fig. 3 d), and continued to grow toward the ovary. After penetrating the ovary, the pollen tube continued to grow along the placenta’s epidermal cells (Fig. 3 e). After 48–60 h of flowering, when the pollen tube approached the ovule, it was bent nearly 90° to approach and penetrate the micropyle and pass through the nucellus into the embryo sac (Fig. 3 f). A few aborted pollen grains could be seen on the stigma after pollination, as well as some abnormal pollen tubes and self-pollinating pollen tubes. However, due to the large amount of pollen, sufficient pollen tubes entered the ovary, enabling us to observe each ovule in the ovary with the pollen tube entering.

**Fig. 3.**
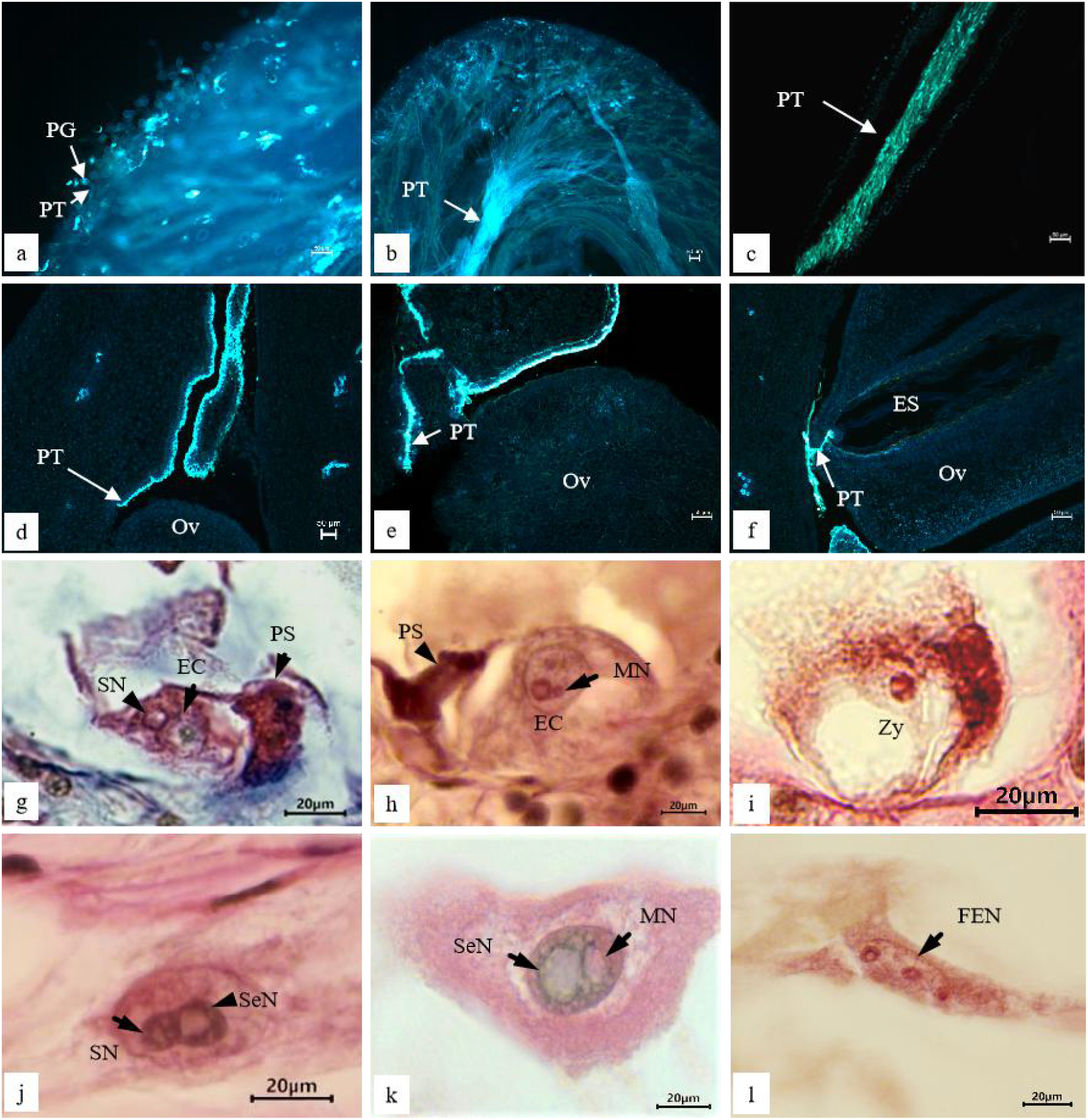
Pollen tube growth and double fertilisation process after natural pollination of *P. ludlowii:* **(a)** pollen tubes germinated on the stigma and grew into the stigma (laminated fluorescence); **(b)** pollen tubes converged into the style and grew downward (laminated fluorescence); **(c)** pollen tubes grew along the stylistic tract to the ovary (section fluorescence); **(d)** the pollen tube entered the ovary along the style channel (section fluorescence); **(e)** pollen tube reached near the funicle of ovule along the placenta (section fluorescence); **(f)** pollen tube entered the embryo sac through the nucellus (section fluorescence); **(g)** the sperm nucleus attached to the egg nucleus; **(h)** the sperm nucleus entered the egg nucleus, showing the male nucleoli (mn) in the egg nucleus; **(i)** zygote formation; **(j)** the spermatic nucleus attached to the secondary nucleus; **(k)** male nucleoli appeared in the polar nucleus; and **(l)** free endosperm nucleus formation. EC, egg cell; ES, embryo sac; FEN, free endosperm nucleus; MN, male nucleus; Ov, ovule; PG, pollen grain; PT, pollen tube; PS, persistent synergid; SN, sperm nucleus; SeN, secondary nucleus; Zy, zygote.

#### Double fertilisation

After passing through the apical nucellus from the degenerated synergid entering the embryo sac, the pollen tube released two sperm cells, one fusing with the egg nucleus and the other with the secondary nucleus (polar nucleus) of the central cell. The sperm nucleus gradually approached the egg nucleus, fused with it, and eventually formed large nucleoli of the zygote, and the fertilisation of the egg cell ended (Fig. 3 g–i). This process can be observed at 60–144 h after flowering. After zygote formation, a period of dormancy was required before the division stage. The fusion of the sperm nucleus and secondary nucleus was similar to the fusion of the sperm nucleus and egg nucleus (Fig. 3 j, k). The difference was that the fertilisation rate of the central cell was significantly higher than that of the egg cell, as determined by observing several successive sections. After fertilisation of the central cell, the primary endosperm nucleus was formed; this nucleus then divided to produce the free endosperm nucleus (Fig. 3 l). This process can be observed from 60 to 108 h after flowering. The timing of double fertilisation of ovules in each ovary was not synchronous. During 108–144 h after flowering, some ovules in the double fertilisation stage were also observed, but the number was small; the process was completed by day 7 after flowering.

During the late development of fertile ovules in *P. ludlowii*, the primary endosperm nucleus was the first to change. After the fertilisation of the secondary nucleus (polar nucleus), the primary endosperm nucleus split to form several free endosperm nuclei inside the embryo sac (Fig. 4 a). Subsequently, the free endosperm nuclei were divided repeatedly, and the number of free nuclei increased continuously, during which no cell wall was formed (Fig. 4 b–d). Until 45 d after flowering, the free nuclear endosperm began to cellularise, and at this time, the internal ovule was liquid or semi-liquid (Fig. 4 e). The cellularisation was completed at approximately 55 d after flowering (Fig. 4 f). Thereafter, the volume of the endosperm increased rapidly and the endosperm reached its final shape and size approximately 75 d after flowering and the interior of the ovule gradually changed from liquid or semi-liquid to a solid state.

**Fig. 4.**
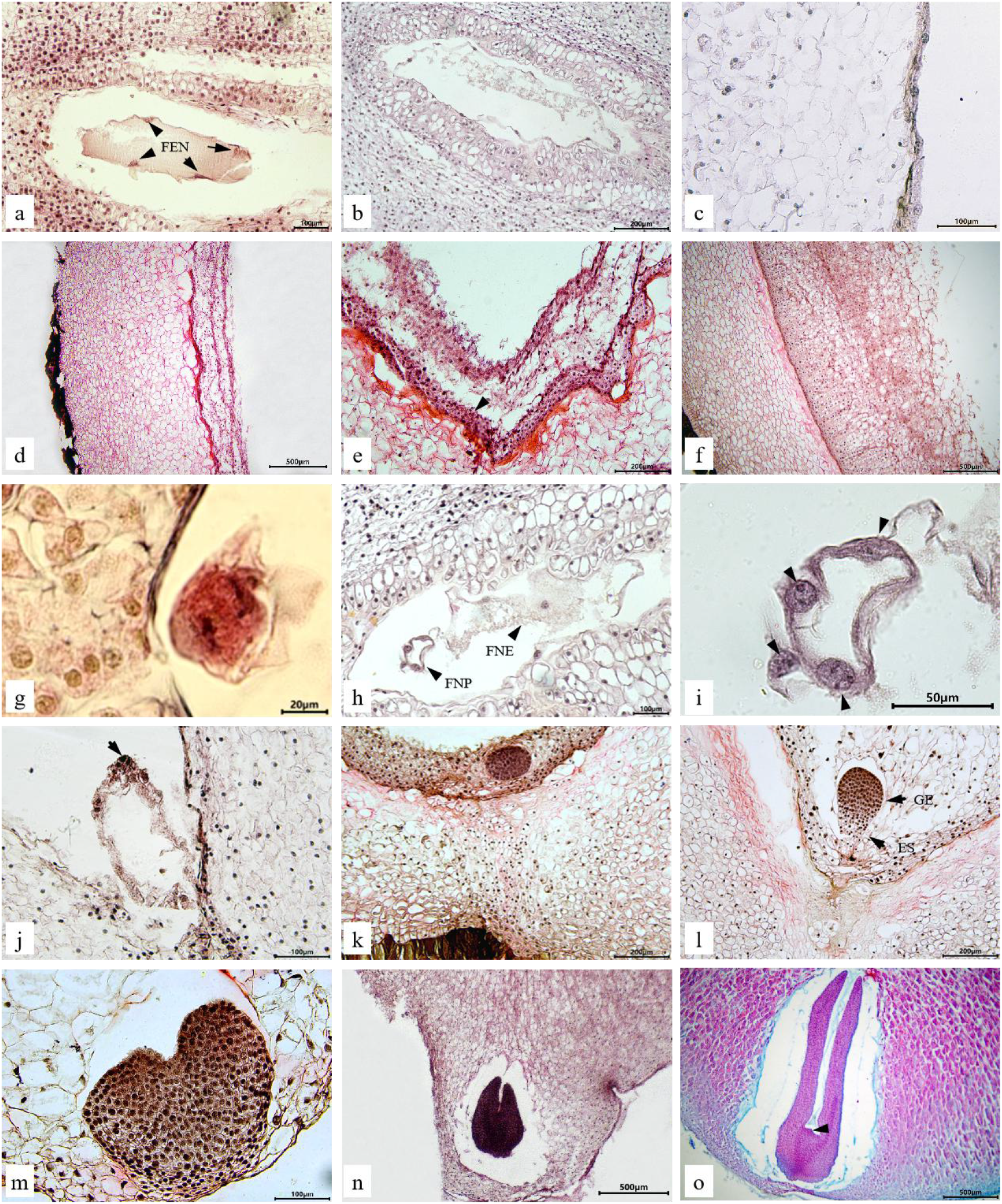
Development of the endosperm and embryo in a fertile ovule of *Paeonia ludlowii*: (**a–c**) at 120 h, 12 d, and 20 d after flowering, the free nuclei of the embryo sac and endosperm continued to increase, the nucellus and inner tegmentum degenerated, and the endosperm adjoined the outer tegmentum; (**d**) the free nuclei of the endosperm continued to increase, and the degraded area of the pearly was formed (40 d after flowering); (**e**) 45 d after flowering, the endosperm began to cellularise (arrow shows the cellularised endosperm); (**f**) 55 d after flowering, the endosperm was cellularised; (**g**) telophase of the first zygote division (108 h after flowering); (**h, i**) the embryo sac of free nuclear proembryo and free nuclear endosperm (12 d after flowering), (**i**) shows the amplification of free nuclear proembryo in h (arrow shows free nuclei); (**j**) the free nuclei at the chalazal end began to cellularise (30 d after flowering) (arrow shows the cellularised embryo); (**k**) early stage of spherical embryo (55 d after flowering); (**l**) late globular embryo (60 d after flowering); (**m**) heart-shaped embryo (65 d after flowering); (**n**) torpedo embryo (70 d after flowering); (**o**) cotyledons at seed maturity (arrow indicates points of growth) (100 d after flowering). FNN, free nuclear endosperm; FNP, free nuclear proembryo; GE, globular embryo; ES, embryo stalk.

The endozygote of the fertile ovules began to divide after the end of dormancy (Fig. 4 g). The zygote divided first to form the binuclear proembryo and then divided repeatedly and synchronously. The number of free nuclei increased continuously, and the proembryo grew gradually. During this period, no cell wall was formed (Fig. 4 h, i). The free nuclear stage of the proembryo lasted from zygotic dormancy to 30 d after flowering. The free nuclei then began to cellularise, generally beginning at the chalazal end and advancing towards the micropylar end (Fig. 4 j). At 55 d after flowering, embryo development reached the globular embryo stage (Fig. 4 k, l). Thereafter, the embryonic somatic cells divided and differentiated rapidly, and the embryonic development successively went through the heart-shaped embryo stage (Fig. 4 m), torpedo-shaped embryo stage (Fig. 4 n), and cotyledon embryo stage (Fig. 4 o), until the final seeds were ripe.

### Abnormalities during fertilisation and seed development

#### Abnormal pistil

During the growth and development of *P. ludlowii,* some pistils were abnormal in different ways, for example, ovule bursting, caused by the rupture of the carpel (Fig. 5 a), stigma exposure before anthesis (Fig. 5 b), and carpel deformity (Fig. 5 c–f).

**Fig. 5.**
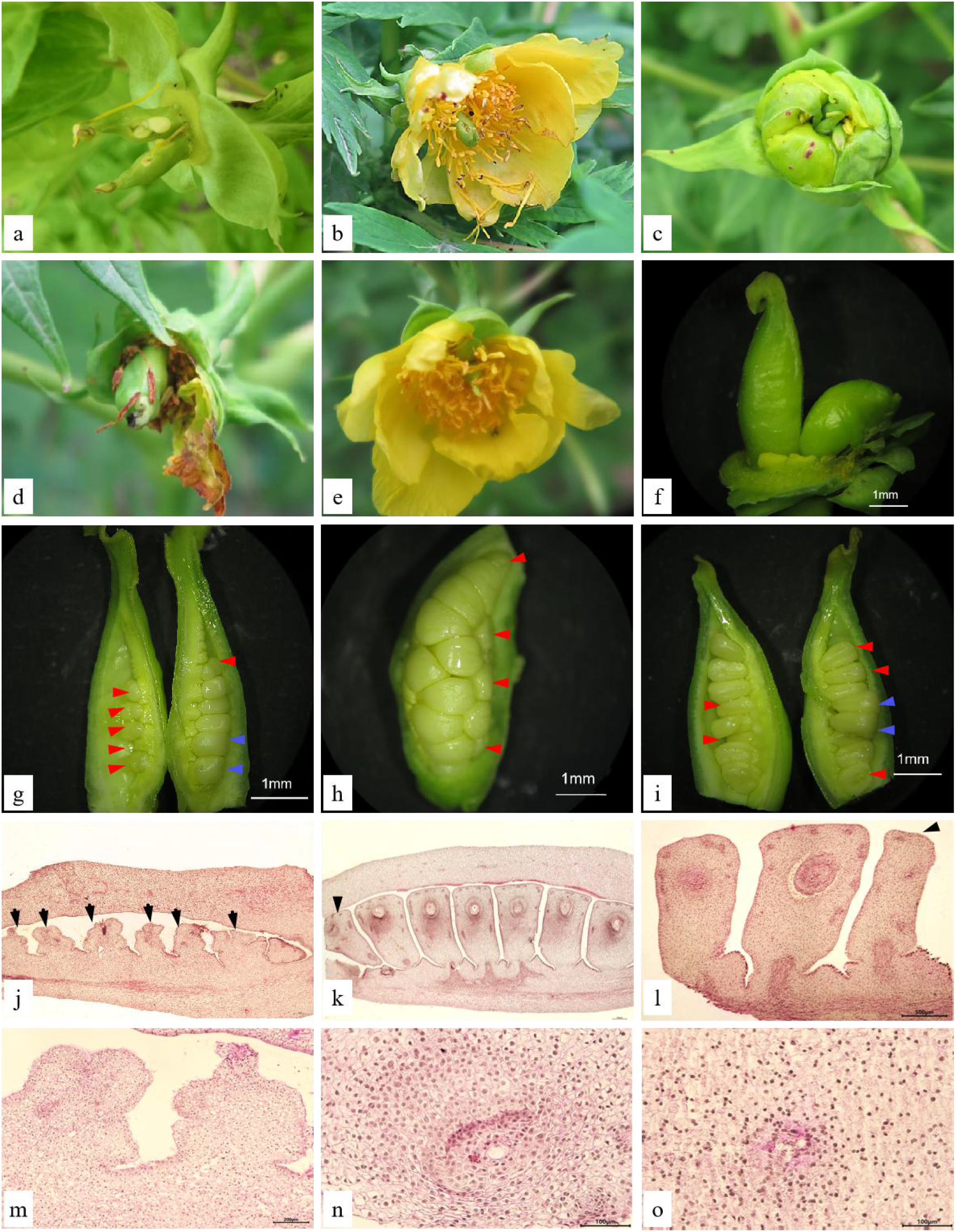
Abnormal pistils and sterile ovules of *P. ludlowii:* **(a)** the ovary was ruptured, and the ovules could be seen protruding; **(b)** stigma showing **(c)** abnormally curved carpel; **(d)** abnormal style and stigma; **(e, f)** one carpel of flower without stigma or with a regressed stigma; **(g–i)** sterile ovules at different position in the ovary (sterile and fertile ovules are indicated by red and blue arrows); and **(j–o)** microstructure of abortive ovules in the ovary, (**j**) and (**m**) show that the whole row of ovules in the ovary were aborted (on the first day of flowering), (**k**) and (**n**) show the ovules in a row that were aborted in the upper position of the ovary, and (**l**) and (**o**) indicate ovule abortion in the middle position.

These pistils were present before flowering and could not be fertilised normally. Among nearly 500 flowering branches and about 1,500 open flowers (some buds failed to open normally), about 30 had abnormal pistils, accounting for 0.02%.

#### Sterile ovule

During the study, repeated experiments and many dissections showed that the appearance of carpels in the ovaries of some test materials was the same as that of other carpels, but there were sterile ovules inside. These ovules were present on the day of flowering, when the embryo sac matured, rather than being abnormal during post-pollination development. These ovules were markedly different from other ovules in morphology; that is, they were smaller in size or deformed (Fig. 5 g, h, i). Some of them had no embryo sac, and there were no clear spaces between the outer and inner integument (Fig. 5 j, m). Moreover, some of the ovules had embryo sac structures, and the space was small, without any obvious organisational structure (Fig. 5 k, l, n, o). In the materials observed, not all carpels had sterile ovules, and according to the statistics, the carpels with sterile ovules accounted for 10.59%–15.73% of all the test materials.

#### Abnormal embryo sac

Through several sections and repeated experiments, we found some abnormal embryo sacs in the experimental materials during fertilisation. The main characteristics of these abnormal embryo sacs were as follows. 1) At 84 h after anthesis, the pollen tube had reached the ovary, and most ovules had begun fertilisation, but the two synergistic cells in the individual embryo sac remained intact without degeneration. The embryo sac, which could not enter the physiological state of fertilisation in time, could not be fertilised normally (Cheng, 1996). 2) At 8–12 d after flowering, free nuclear proembryo and endosperm were formed in the fertilised ovules, and the number of free nuclei increased significantly, but the secondary nuclei in some of the embryo sacs remained unfertilised, and no endosperm free nuclei were formed. These ovules could not develop into seeds normally without double fertilisation. The ovules containing abnormal embryo sacs accounted for about 1.2% of all experimental materials, and only a few carpels had abnormal embryo sacs.

#### Degeneration of the embryo and endosperm

There were some ovules in the carpel, which could complete double fertilisation, and showed zygote and primary endosperm nuclear division in the early stages but could not develop further in the late stages. At 9 d after flowering, some aborted ovules similar in size to normal ovules showed abnormalities in the embryo sac, and the free nuclei of the endosperm had degenerated (Fig. 6 a, b). On day 10 after flowering, the free nuclei of the endosperm continued to degenerate (Fig. 6 c). Some abortive ovules of the same size as normal ovules showed shrinkage and deformity in their outer tepals after dewatering and embedding (Fig. 6 d). When no free nuclear embryo was observed in the embryo sac, the endosperm developed abnormally, and the free endosperm nuclei were rare and irregular. The cytoplasm of the endosperm gradually disintegrated and was in a state of severe degeneration (Fig. 6 e, f). In the material at 11 d after flowering, apart from the phenomenon of endosperm abortion, the free endosperms in some of the embryo sacs aggregated after nuclear division, but did not separate (Fig. 6 g). At 12 d after flowering, the endosperm of the fertilised ovules degenerated. In some embryo sacs, only the free nuclear proembryo developed to the stage of dinucleus proembryo, but the free endosperm nuclei were rare, the cytoplasm was significantly reduced, and the embryo sac was in a severely degenerated state; consequently, the embryo sac became narrow (Fig. 6 h). Embryo sacs were distinct from the fertile ovules at the same period (Fig. 4 h). On the same day, with the development of the ovaries and the enlargement of the ovules, the ovules’ volume in the ovaries began to show differences (Fig. 6 i). A few aborted ovules developed normally until approximately 20 d after flowering, and the volume of the ovules was slightly larger than that of other early aborted ovules and slightly smaller than that of fertile ovules (Fig. 6 j, indicated with the arrows). The free nuclei of the embryo and endosperm in these aborted ovules were not as developed as the other fertile ovules and tended to degenerate (Fig. 6 k). At this time, less than one-third of the fertile ovules were observed in the ovary, and abortive ovules were observed in all carpels. The results indicated that the abortion of embryo and endosperm occurred mainly in the early stage, 9–12 d after flowering.

**Fig. 6.**
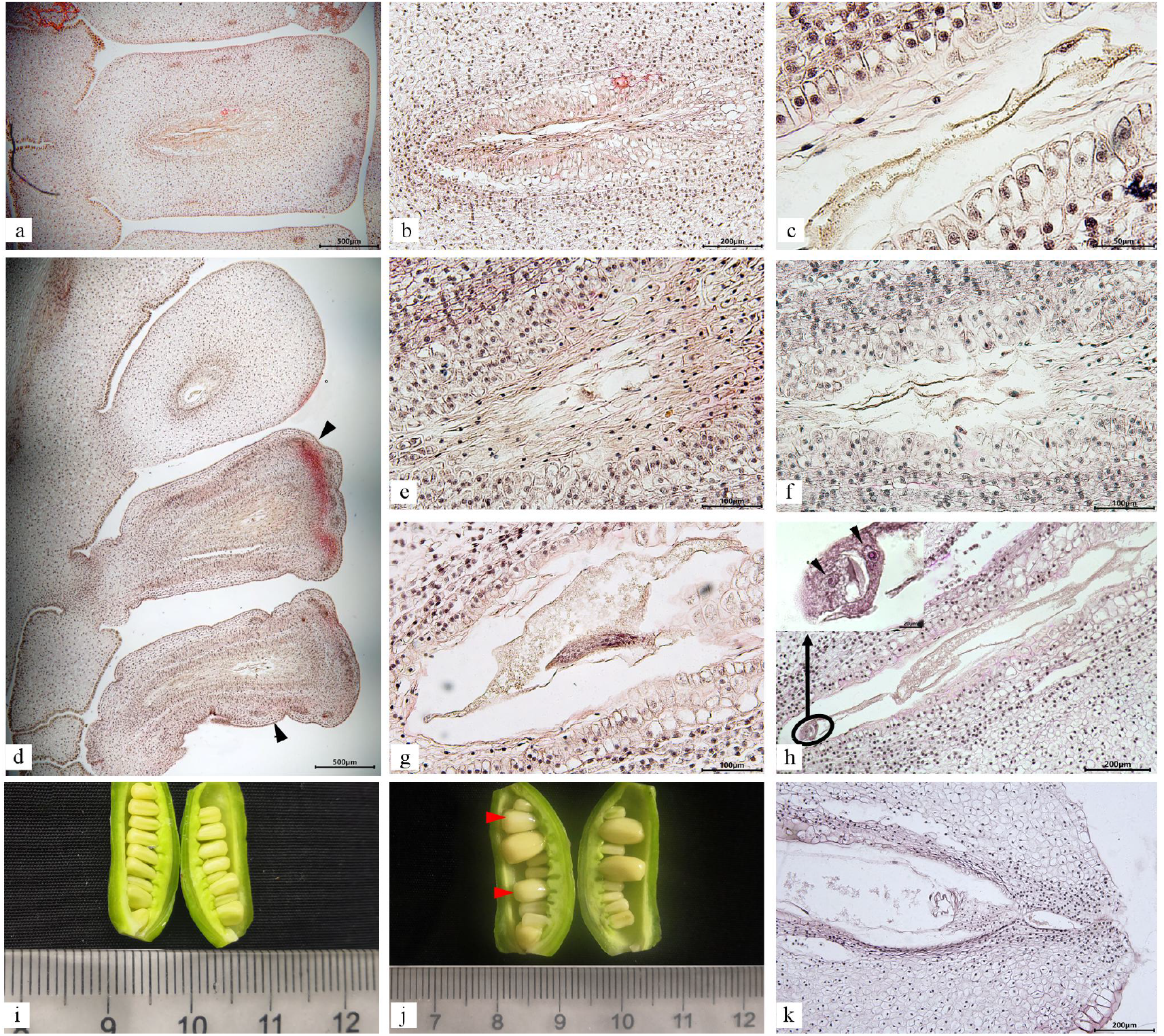
Degradation of embryo and endosperm after double fertilisation of aborted ovules in *Paeonia ludlowii:* **(a, b)** the endosperm in the embryo sac was degenerative. The cavity of the embryo sac became narrow 9 d after flowering; **(c)** aborted ovules at 10 d after flowering; **(d, e, f)** two successive aborted ovules (10 d after flowering), (**e**) is the enlargement of the embryo sac in d-①, (**f**) is the enlargement of the embryo sac in d-②, the cytoplasm of the endosperm was reduced, and most of the free nuclei had degenerated; **(g)** there was free endosperm nuclear division, but no separation (11 d after flowering); **(h)** the embryo in the embryo sac was in the dikaryotic proembryo stage, but the free nucleus of the endosperm had degenerated, only a part of the cytoplasm remained, and the embryo sac became small (12 d after flowering); (**i**) at 12 d after flowering, the volume of the ovules in ovaries was different; **(j, k)** at 20 d after flowering, a part of normally developing ovules in the early stage was aborted (**k** the internal structure of the embryo sac shown with the arrow in **j**).

#### Late development of abortive ovules

The development of early aborted ovules stagnated at 12 d after flowering, and the volume of aborted ovules was significantly smaller than that of the normal ovules (Fig. 6 i). After 12 d, the embryo sacs of some aborted ovules gradually contracted towards the nucellus, the cavity of the embryo sac became narrow and long, degenerated, and necrotic, and filled with brown material (Fig. 7 a, b). At 30 d after flowering, the embryo, nucellar, and integument gradually degenerated (Fig. 7 c). Some embryo sacs were filled with large cells, with only a narrow gap in the middle, with darker staining (Fig. 7 d–g). With the growth of the ovule, the free nuclear embryo and endosperm of some ovules that began to abort at approximately 20 d after flowering degenerated completely, and the outer envelope had also gradually degenerated, finally forming a hollow shell with no embryo or endosperm in the embryo sac. The outer cover gradually degenerated, finally forming a hollow shell without an embryo and endosperm in the embryo sac (Fig. 7 h–j). Although the embryo and endosperm were not formed in the abortion ovules, the external morphology of the abortion seeds significantly differed (Fig. 7 k, shown by the red arrow). In addition, when harvesting seeds, we also found a few seeds with external morphology and volume similar to those of the fertile seeds, but they were deformed after applying pressure. After peeling, the seeds were hollow inside, without embryo or endosperm structure (Fig. 6 l).

**Fig.7.**
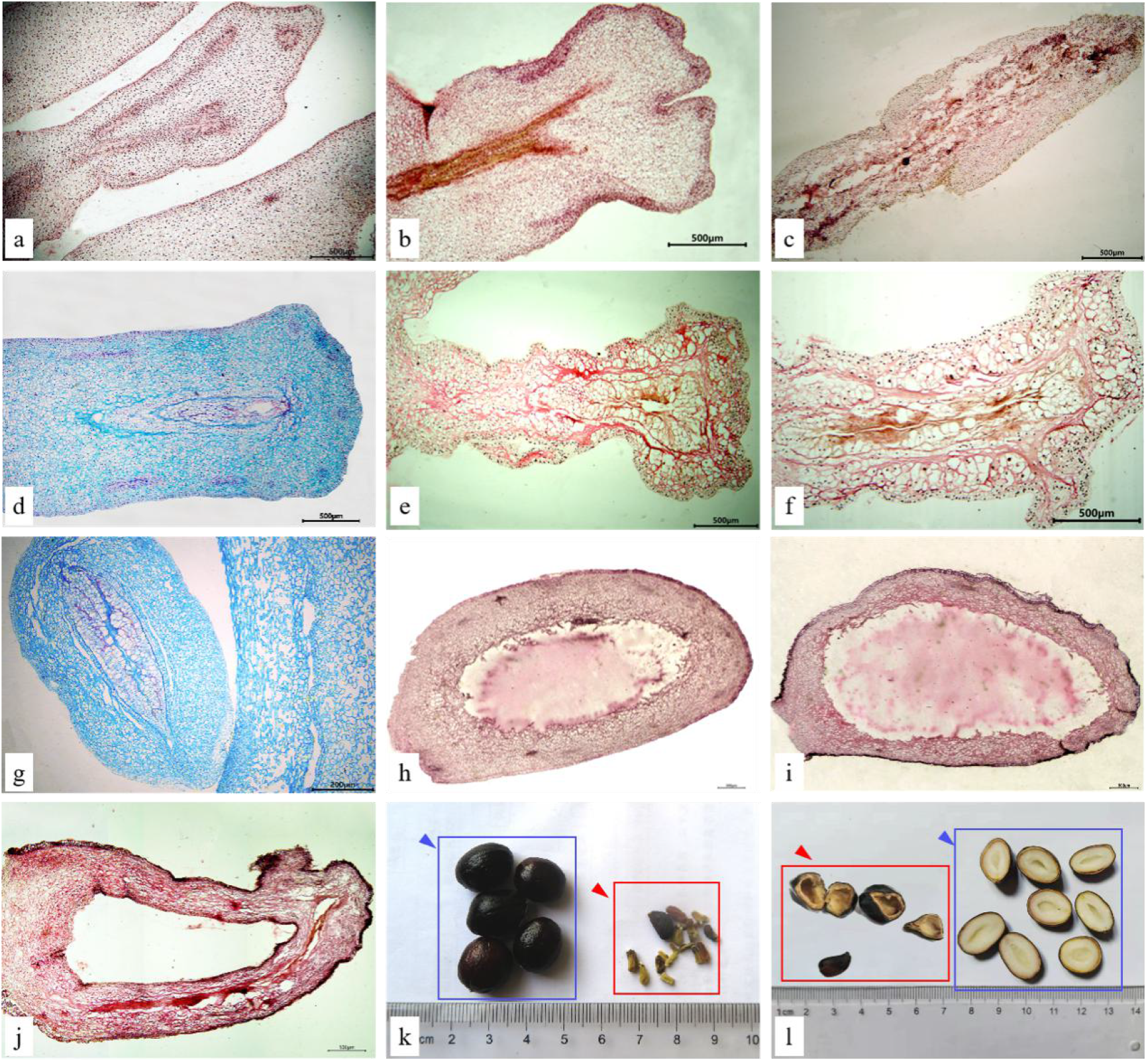
Late development of aborted ovules: **(a–d)** the development of aborted ovules at 12, 25, 30, and 35 d after flowering; **(e, f)** development of aborted ovules at 55 d after flowering; **(g)** development of aborted ovules at 70 d after flowering; **(h–j)** development of the aborted ovules of free nuclear embryos and endosperm aborted at 20 d after flowering at 30, 45, and 70 d after flowering, respectively; **(k)** comparison of fertile ovules and aborted ovules at seed harvest (aborted and normal seeds are indicated with red and blue arrows, respectively); and **(l)** comparison of the internal structure of aborted ovules and fertile ovules at seed harvest (aborted and normal seeds are indicated with red and blue arrows, respectively).

### Position and rate of aborted ovules in *P. ludlowii*

The total number of seeds in a single carpel was 6–16, and the number of normal seeds was 0–8 (2–4 more). More than two-thirds of the ovules were aborted (Table 1, Fig.8). According to the statistics of the rate of abortion seeds (Table 1), the seed abortion rate in the ovaries in 2019 and 2020 was 69.14% and 74.01%, respectively, and the abortion rate in wild populations was also high (66.39%), indicating that seed abortion occurred independent of the growing environment.

**Table 1.** Ovule position and abortion rate in *Paeonia ludlowii*.

| Year | Number of seeds in a single ovary |  |  | Position and rate (%) of ovule abortion |  |  | Mean (%) |
| --- | --- | --- | --- | --- | --- | --- | --- |
|  | Total number of seeds | Normal seeds (mature seeds) | Abortion seeds | Upper part of the ovary | Middle part of the ovary | Lower part of the ovary |  |
| 2019 | 6–16 | 0–8 | 4–14 | 66.25 ± 3.20b | 66.98 ± 2.66b | 77.08 ± 1.65a | 69.14 ± 1.41a |
| 2020 | 6–16 | 1–8 | 4–14 | 70.86 ± 2.813b | 83.33 ± 2.484a | 78.59 ± 1.523a | 74.01 ± 1.06b |

Note: Different lowercase letters indicate significant difference at the 0.05 level ( $P < 0.05$ ).

**Fig. 8.**
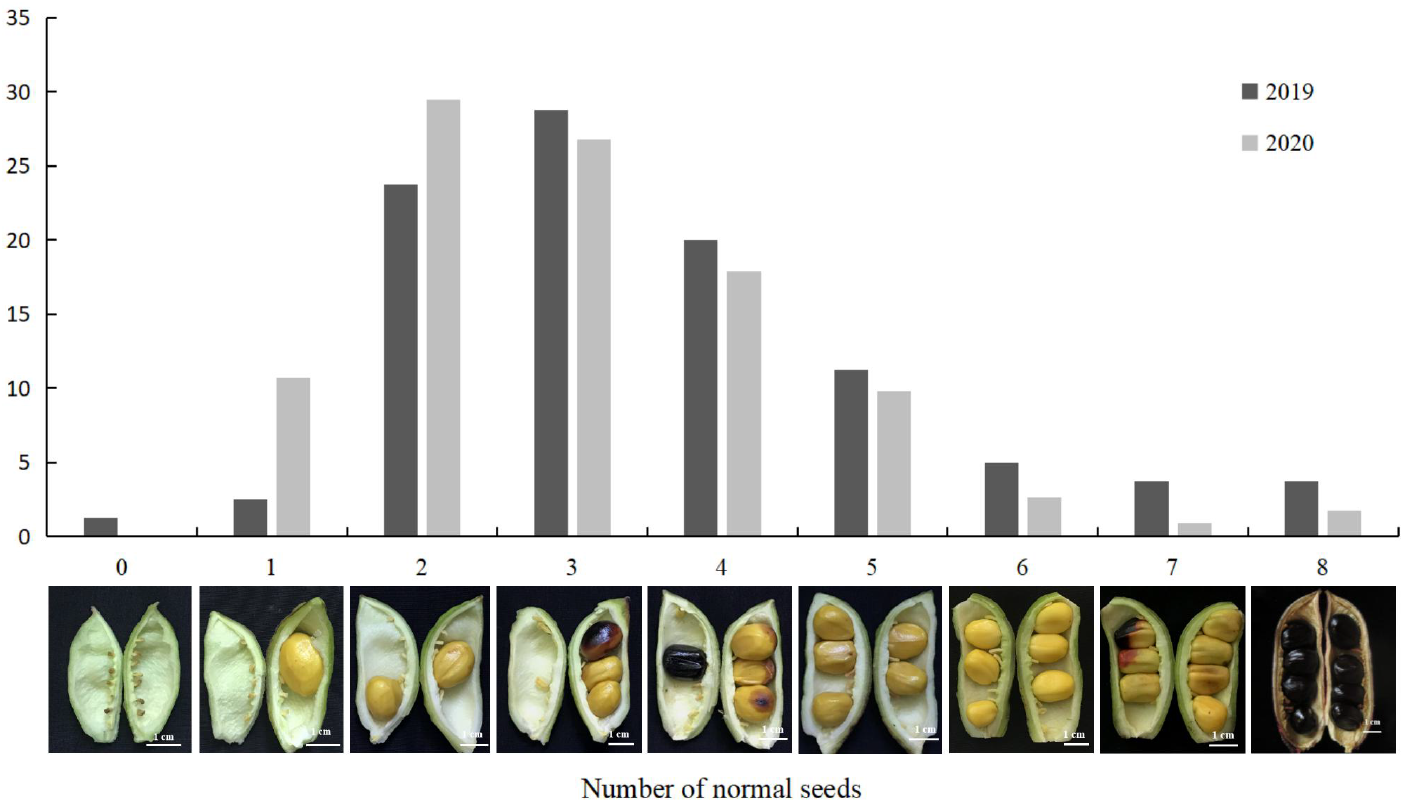
Distribution of the number of normal seeds in *Paeonia ludlowii* ovaries.

The statistical results of the position of abortion seeds in *P. ludlowii* (Table 1) showed that, in 2019, the abortion rate in the lower part of the ovary was the highest (77.08%) and the abortion rates in the middle and upper parts of the ovary were 66.98% and 66.25%, respectively. Moreover, the difference among the three parts was significant. In 2020, the abortion rate was the highest in the middle part of the ovary (83.33%), followed by the lower part (78.59%) and the upper part (70.86%). These results showed that there was no specific position for seed abortion in the ovary of *P. ludlowii,* and its seed abortion was random.

## Discussion

### Characteristics of the carpel and embryo sac of *P. ludlowii*

The carpels are plant reproductive organs, and ovary development (ovule development and embryo sac formation) and stigma and style formation are particularly important for fertilisation (Bedinger et al., 2017; Gao et al., 2019). The carpel of tree peony is composed of the ovary, style, and stigma. Zhao et al. (2002) found that the stigma of *P. ludlowii* curled at an angle of about 90°. However, in this study, most of the stigmas curled at an angle of 360° or nearly 360° into a ring. The results of this study were slightly different from those of previous studies; this could be because, in order to adapt to the environment and to meet the needs of pollination, the stigma had undergone a certain degree of evolution, which enlarged the pollen surface. In addition, the electron microscopy structure of the stigma showed that the stigma surface of *P. ludlowii* is covered with a large number of papillae, which also helps the stigma to accept many pollen grains.

The style of *P. ludlowii* is hollow, with a canal in the centre. As in the lily and camellia, which have hollow styles, the channel is surrounded by a layer of glandular secretory cells, also known as channel cells (Hu et al., 1982; Gao et al., 2019; Zhang, 2020). The style is the only way that the pollen tubes can enter the ovary. In plants with style channels, pollen tubes grow mainly along the surface of the channel cells and then enter the ovary (Gao et al., 2015a).

The ovules of *P. ludlowii* are inverted, with a double integument, thick nucellus, and polygonum-type embryo sac. The basic structure is the same as that of other peonies (Cheng and Aoki, 1999; Wang et al., 2010). The embryo sac of *P. ludlowii* developed and matured earlier, with one synergistic cell degenerating at 12 h after flowering, and the embryo sac entered the fertilisation state. Walters’ (1962) research on *P. californica* revealed that each ovule had 1–4 embryo sacs, and the phenomenon of multiple embryo sac developmnt has also been found in the ovules of *P. rockii* (Cheng and Aoki, 1999). However, only one embryo sac was observed for each ovule in many materials of *P. ludlowii,* this is consistent with the finding of Wang et al. (2010) and Vinogradova et al.(2020), that is, only one embryo sac is formed in *P. lactiflora* species.These results indicate that embryo sac development in *P. ludlowii* was normal and that it had strong sexual reproduction ability.

### Characteristics of pollen tube growth in *P. ludlowii*

The completion time of pollen grain germination and pollen tube growth is 2–3 h in *P. rockii* (Cheng, 1996). However, the pollen grains of *P. ostii* ‘Feng Dan’ germinate immediately after falling on the stigma and form pollen tube channels 1 h after pollination, and the pollen tubes enter the ovule and reach the embryo sac at 12–48 h after pollination (Fan et al., 2004; Dong, 2010; Chen, 2020). In ‘High Noon’ (a distant hybrid between subgroups in the peony group) (Walters, 1942), germination is not observed until 8–11 d after pollination, when the pollen tube enters the nucellus (He and Cheng, 2006). Here, the observation results for *P. ludlowii* were similar to those of *P. ostii* ‘Feng Dan’ and the pollen tubes in *P. ludlowii* entered the embryo sac at 48–60 h after flowering. Pollen germination capacity and pollen tube growth rate in plant styles are affected by internal and external factors, such as the variety characteristics, nutritional conditions, temperature, light, and climate, as reported in some species and varieties (Schlichting, 1986; Lau and Stephenson, 1994; Dogterom et al., 2000; Ruane, 2008; Tuell and Isaacs, 2010). Thus, the germinated pollen tube takes a longer time to enter the embryo sac in *P. ludlowii* than in *P. rockii* due to varietal differences, planting conditions, and climatic conditions after pollination.

In angiosperms, the pollen tube enters the ovule in two ways namely porogamy and chalazogamy. When the pollen tube of *P. ludlowii* approached the ovule, it turned nearly 90° and entered the micropyle. Therefore, the fertilisation pattern of *P. ludlowii* is porogamy, as in *Chrysanthemum grandiflorum* and *Camellia oleifera* (Deng et al., 2010; Gao et al., 2015b). The double fertilisation of *P. ludlowii* is consistent with that in *P. rockii* (Cheng, 1996), indicating that double fertilisation in *P. ludlowii* is normal. According to a report by Dong Zhaolei (2010), in *P. ostii* ‘Feng Dan’, two sperms enter the egg cell at the same time or near the secondary nucleus. However, we did not observe this phenomenon in *P. ludlowii.* Whether it was related to the experimental materials remains to be studied.

### Characteristics of double fertilisation and development of embryo and endosperm in *P. ludlowii*

The development of endoembryo and endosperm in the fertile ovule of *P. ludlowii* is the same as that in other tree peony species (Cave et al., 1961; Mu and Wang, 1995; Cheng and Aoki, 1999; Dong, 2010), but the timing is significantly different (Table 2). This difference can be attributed to the plant species and environmental conditions, such as flowering and early embryo development temperatures.

**Table. 2.** Initiation time of free nuclear proembryos and endosperm in *Paeonia*.

| Plant species | Growing environment (location) | Cellular time of proembryo (days after flowering or pollination, d) | Cellular time of endosperm (days after flowering or pollination, d) | Source |
| --- | --- | --- | --- | --- |
| *P. californica*<br>*P. brownii* | Wild (California, US) | > 12–14 | > 12–14 | Cave et al. (1961) |
| *P. lactiflora* | Cultivated (Beijing, China) | 17–23 | > 17–23 | Xijin Mu et al. (1985) |
| *P. rockii* | Cultivated (Lanzhou, China) | 29 | 29 | Fangyun Chen (1999) |
| *P. ostii* ‘Feng Dan’ | Cultivated (Shanghai, China) | 23 | 23 | Zhaolei Dong (2010) |
| *P. ludlowii* | Cultivated (Luoyang, China) | 30 | 45 | This study |

### Ovule abortion and its characteristics in *P. ludlowii*

In seed plants, abortion could be divided into nonrandom and random abortion. The former refers to a certain regularity in the position of seed abortion in the fruit, as in *Phaseolus coccineus* (Rocha et al., 1990), *Robinia pseudoacacia* (Susko, 2006; Yuan et al., 2014), *Caesalpinia gilliesii* (Calviño, 2014), *Bauhinia ungulata* (Mena-Ali et al., 2004), and *Anagyris foetida* (Valtueña et al., 2010). The latter means that the abortion seeds have no specific position in the fruit. Evidently, the abortion of *P. ludlowii* belongs to the latter.

The aborted ovules of *P. ludlowii* are of four types: abnormal pistils, sterile ovules, abnormal embryo sacs, and embryo and endosperm abortions. The phenomenon of pistil abortion, such as abnormal pistils, sterile ovules, and abnormal embryo sacs, is common in nature (Wang, 2006; Wetzstein et al., 2011; Hou et al., 2011). However, although pistil abortion exists, ovule abortion in *P. ludlowii* mainly involves embryo and endosperm abortions. According to the time, embryo and endosperm abortions in *P. ludlowii* could be divided into three types. 1) Early abortion, which occurred at 9–12 d after flowering and the aborted ovules stopped growing and were smaller than fertile ovules. The development of endosperm was abnormal, the free nucleus of the endosperm continued to degenerate, and the embryo sac became narrow. The number of abortive ovules in the ovary was more than 60%, and all carpels had abortive ovules. 2) Medium-term abortive, which occurred about 20 d after flowering. The volume of aborted ovules was slightly larger than that of other early aborted ovules and slightly smaller than that of fertile ovules. The free nuclei of the embryo and endosperm in the embryo sac began to degenerate. 3) Late abortive, wherein the abortion seeds were similar in external morphology and size to the fertile seeds, but after being peeled, the inside was hollow, without an embryo and endosperm structure. There were only a few seeds of this type. Based on the proportion of abortion ovules observed in each stage, the embryo and endosperm of *P. ludlowii* were mainly aborted in the early stage, that is, 9–14 d after flowering. The occurrence time was slightly different from the results of He et al. (2006), which might be related to various characteristics of the plant and different pollination and fertilisation times.

Several factors besides species characteristics and resource constraints affect plant embryo abortion (Nakamura, 1988; Teixeira et al., 2006;Brookes et al., 2008; Florez-Rueda et al., 2016). These include sibling competition (Ganeshaiah et al., 1988; Jerry et al., 2015), poor pollination and fertilisation (Daiichiro et al., 2003; Shen et al., 2018; Xie et al., 2019), endosperm abortion (Sun et al., 2009; Pan et al., 2011;), abnormal hormone metabolism (Okamoto et al., 1991; DeBruin et al., 2018; Daniela et al., 2020a; Daniela et al., 2020b), and nutritional supply imbalance (Ji et al., 2019; Shen et al., 2020). In this study, there was no abnormality in the process of pollination and fertilisation, so the factor of poor pollination and fertilisation could be excluded. During flowering and fruiting, the axillary buds on the flowering branches of *P. ludlowii* sprouted into the secondary branches, and the apical buds on the secondary branches differentiated (Yuan et al., 2021). Thus, the fruit development period coincided with the secondary branch growth period and the apical bud differentiation period. Consequently, the nutrients should be balanced and distributed among the three, and this may be one of the causes of ovule abortion. In addition, during the early and middle stages of embryo and endosperm abortion, the endosperm degenerated first, and the embryo degenerated later, which might be because the endosperm acts as an important source of nutrients for embryo development and can store nutrients such as starch, lipids, and proteins (Lopes et al., 1993; Gehring, 2004). When the endosperm degenerated, the embryo lacked nutrients and was aborted. In conclusion, the reasons for ovule abortion in *P. ludlowii* are complex and influenced by several factors. The findings of this study expand our understanding of the source and characteristics of ovule abortion, and the specific factors of ovule abortion should be an important research direction in the future.

## Materials and Methods

### Materials

The materials used in this study were collected from the ex-situ Conservation Center of Chinese Paeoniaceae Wild Species (Little Red Village, Sanchuan Town, Luanchuan County, Luoyang City, Henan Province, China) (111°21’36”E, 33°56’05”N; 1,245 m altitude).*Paeonia ludlowii* plants were sown in the autumn of 2002 (the seeds had been collected from the wild population). Currently, the plants have been growing vigorously, and flowering and fruiting have been stable, with nearly 1,500 flowers blooming every year. Light management has been applied, with weeding once a year.

### Methods

The experiment was conducted from May to September 2019 and from May to September 2020. The experiment was repeated twice for 2 consecutive years.

### Unfertilised carpel and embryo sac were observed by paraffin sectioning

The carpels of *P. ludlowii* were collected before the flowers had bloomed. After collection, the stigma, style, and ovary were cut off. The styles and ovaries were placed in Carnoy’s solution (95% ethanol:glacial acetic acid [v/v], 3:1). After 12 h of vacuum treatment, the materials were transferred to 70% ethanol solution and stored in a refrigerator at 4 °C. The stigmas were then placed in a fixative (50% ethanol:acetic acid:formaldehyde [V/V/V] = 90:5:5). After 48 h of vacuum treatment, the materials were washed twice in 50% ethanol solution, each time for 2 h, and then transferred to 70% ethanol and stored in a refrigerator at 4 °C. After proper trimming, the fixed style and ovary samples were embedded in conventional paraffin sections, and the samples were sectioned using a microtome (RM2235; Leica, Germany) to a thickness of 8 μm. The sections were stained with haematoxylin, and permanent preparations were generated using Canada balsam. Images were captured under a microscope (CX40-RFL; SDPTOP, Ningbo, China).

### Stigmas were observed by SEM

The fixed stigma samples were transferred to ethanol solution of different concentrations (70%, 80%, 90%, 95%, and 100%) for gradient dehydration and soaked for 15 min at each concentration. The materials were transferred to tertiary butyl alcohol, and then subjected to cryodesiccation. The samples were placed in an ion sputter coater and gilded for 20 min. The stigmas were observed by SEM (S-3400N; Hitachi, Japan).

### Pollen tube growth

The carpels of *P. ludlowii* were collected at 1, 3, 5, 8, 12, 24, 36, 48, 60, 72, 84, 96, 108, and 120 h after the flowers had bloomed. The carpel samples were fixed in Carnoy’s solution (95% ethanol:glacial acetic acid [v/v], 3:1) for 12 h of vacuum treatment, and then stored in 70% alcohol at 4 °C in a refrigerator.

The growth of the pollen tube through the stigma and style was observed using the aniline blue compression method (Kho and Baër, 1968; Cheng et al., 2015). The collected carpel samples were dissected from the middle along the dorsal suture and the abdominal suture and divided into two parts. The carpel samples were then washed in distilled water and transferred into 8 mg L^−^ NaOH solution for 2 h. Thereafter, the samples were rinsed in buffer solution of pH 6.7 (1 mol L^−1^ NaOH solution mixed with 45% glacial acetic acid; pH adjusted to 6.7) for 20 min, and then dyed with 0.1% water-soluble aniline blue dye for 6–10 h. The carpel samples were carefully taken out and spread on the slide. A drop of aniline blue solution was added, and the sample was covered with a cover glass and gently pressed. The growth of the pollen tube in the stigma and style were observed, and photographs were taken using a fluorescence microscope (DM2500; Leica).

The growth of the pollen tube in the style, ovary, and ovule was observed by section fluorescence, referring to the method of Gao et al. (2015b), with slight modifications. The specific methods were as follows. According to the pollen tube fluorescence observations in the early stage, the carpel samples after the pollen had reached the style tract were selected and then trimmed and embedded in conventional paraffin sections. After proper trimming, the fixed style and ovary samples were embedded according to the conventional paraffin section method mentioned above. The samples were sectioned using a microtome (RM2235; Leica) to a thickness of 14 μm. After dewaxing, the carpels were soaked in pH 6.7 buffer for 20 min, stained with 0.1% aniline blue solution for 2 h, and then observed and photographed under a fluorescence microscope (DM2500; Leica).

### Double fertilisation and seed development were observed by paraffin sectioning

The carpels at 48 h, 60 h, 72 h, 84 h, 96 h, 108 h, 120 h, 148 h, 168 h, 8 d, 9 d, 10 d, 11 d, 12 d, 13 d, 15 d, 20 d, 25 d, 30 d, 35 d, 40 d, 45 d, 50 d, 55 d, 60 d, 65 d, 70 d, 75 d, 80 d, 90 d, and 100 d after flower blooming were collected. After proper trimming, the carpels were fixed, embedded, and sectioned, following the conventional paraffin sectioning method mentioned above (slice thickness: 9–12 μm). The sections were stained with haematoxylin or saffron-solid green, and permanent preparations were generated using Canada balsam. Images were captured under a microscope (CX40-RFL; SDPTOP).

### Abortion location and abortion rate statistics

When the seeds of *P. ludlowii* were mature, 60 fruits were randomly selected to determine the number of normal seeds (mature seeds), number of abortion seeds, and abortion positions in the ovary. The abortion positions were divided into upper, middle, and lower. The lower part was connected to the fruits, the upper part was near the stigma, and the middle part was at the middle. The abortion rate was then calculated using Eq. 1:

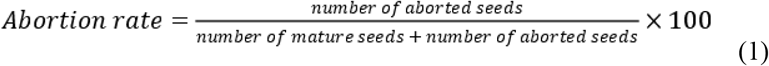

Note: Abortion seeds were undeveloped, flat, empty, brown, and had aborted embryos.

### Statistical analysis

The rate of ovule abortion was analysed using the one-way analysis of variance with IBM SPSS Statistics (version 19.0; IBM Corp., Armonk, NY). A *P* value of <0.05 indicated significant difference. There was extremely significant difference at *P* value of <0.01.

## Acknowledgments

We would like to thank Prof. Fenglan Li and Prof. Jin Cheng for assistance with morphological image identification. We thank Editage (www.editage.cn) for their valuable advice and English language editing.

